# Renal and endothelial biomarkers in Chagas disease in the Brazilian Amazon region: early indicators of kidney injury and disease progression

**DOI:** 10.1101/2025.08.01.668075

**Authors:** Alba Regina Jorge Brandão, Jorge Augusto de Oliveira Guerra, Jessica Vanina Ortiz, Débora Raysa Teixeira de Sousa, Gabriela Maciel Alencar, Nádelly Karoline Martins Derze, João Victor Campelo de Queiroz, Joana Bader Sadala Brandão, Lara Isabelli Oliveira de Silva, Cássia Camila de Oliveira Araújo, Silvia Cassia Brandão Justiniano, Lucely Paiva Rodrigues da Silva, Elsa Isela Moctezuma-Guevara, Susan Smith-Doria, Karla Cristina Silva Petruccelli, Paula Rita Leite da Silva, Mônica Regina Hosannah da Silva e Silva, Kátia do Nascimento Couceiro, Gdayllon Cavalcante Meneses, Letícia Machado de Araújo, Elizabeth de Francesco Daher, Alice Maria Costa Martins, João Marcos Bemfica Barbosa Ferreira, Geraldo Bezerra da Silva Júnior, Maria das Graças Vale Barbosa Guerra

## Abstract

Chagas disease (CD) is a significant public health problem in the Amazon, where the acute phase and oral transmission—often via contaminated açaí—are predominant. These peculiarities influence clinical severity and immunological responses, highlighting the need to investigate non-traditional biomarkers for early detection of renal and endothelial dysfunctions, since conventional methods show limitations in identifying subclinical renal injury. To investigate the expression and clinical relevance of non-traditional biomarkers of renal injury and endothelial activation in acute and chronic CD in the Brazilian Amazon. A cross-sectional study was conducted between July 2021 and July 2023 at the Fundação de Medicina Tropical Dr. Heitor Vieira Dourado (FMT-HVD), Manaus, Amazonas. Seventy-eight native Amazonian patients diagnosed with CD were evaluated and grouped by disease clinical phase: G1a (acute pre-treatment), G1b (acute post-treatment), G2a (chronic indeterminate), and G2b (chronic cardiac). Blood and urine levels of SYN-1, ANG-2, MCP-1, and NGAL were quantified using ELISA and correlated with traditional markers; including creatinine, urea, proteinuria, and glomerular filtration rate (GFR). Biomarkers were elevated across CD phases, despite traditional renal function parameters remaining within normal ranges. Urinary MCP-1 was significantly increased in the acute pre-treatment phase, reflecting early renal inflammation. SYN-1 was elevated in the acute and chronic indeterminate phases, indicating early endothelial activation. NGAL was increased in the chronic indeterminate group and ANG-2 in the chronic phase, suggesting progressive endothelial dysfunction. Traditional markers did not correlate with the biomarkers, suggesting their sensitivity in detecting early subclinical injury. This study highlights the importance of using these biomarkers for early diagnosis and monitoring of CD in the Amazon region. The elevated levels of SYN-1, ANG-2, MCP-1, and NGAL reveal early renal and endothelial dysfunction in CD patients undetectable by conventional tests. Incorporating these biomarkers into routine monitoring could support earlier therapeutic interventions and reduce associated renal and cardiovascular complications.

**AUTHOR SUMMARY:** Chagas disease (CD) is a serious problem in Amazon, where many people get infected by eating contaminated food. If untreated, the disease can silently damage the kidneys and blood vessels. Standard medical tests often miss early signs of the damage, making it hard to intervene before the complications develop. This study aimed to investigate special markers in blood and urine that can indicate early kidney and blood vessel damage in patients at different stages of the disease in the Brazilian Amazon. We discovered that even though standard tests appeared normal, these proposed markers showed signs of damage. MCP-1 was high in early infection, suggesting kidney inflammation; SYN-1 and NGAL were elevated in later stages, suggesting ongoing damage and ANG-2 (linked to blood vessel problems) increased in chronic cases, that could indicate disease progression. In remote areas like the Amazon, where healthcare access is limited, early detection is crucial. The markers described in this study could help doctors detect early, treat and prevent complications like severe kidney and heart damage.

## INTRODUCTION

Chagas disease (CD) is one of the main neglected tropical diseases, with a significant impact on public health, especially in endemic regions of Latin America^1^. This disease can manifest in two clinical phases (acute and chronic) following contact through different routes with the infective form of *Trypanosoma cruzi*^2^. After infection by *T. cruzi*, a cascade of immune responses is triggered, involving inflammation and damage to multiple organs, including the cardiovascular system and the kidneys^3^. In the acute phase of Chagas disease, the parasite multiplies rapidly inside cells, triggering a strong a strong inflammatory response. This leads to intense cellular damage and tissue fibrosis compared to later stages^4,5^. In the chronic phase of patients from areas with classic vector transmission, Chagas disease tends to exhibit greater severity and a higher prevalence of severe chagasic cardiomyopathy, including congestive heart failure and arrhythmias, which are responsible for significant morbidity and mortality^6–8^.

Experimental studies on renal lesions in Chagas disease include functional and structural changes resulting from systemic inflammation, activation of the renin-angiotensin-aldosterone system (RAAS), and possible direct infection of renal tissue by the *T. cruzi* parasite^4,5^. The study of renal involvement in humans with Chagas disease emerges as a relevant aspect of the pathology, due to the interaction between immunological, inflammatory, and autoimmune responses. Furthermore, reduced renal blood flow, activation of the RAAS, and endothelial alterations contribute to the development of renal failure, which is often silent and underdiagnosed^9,10,11^. Traditional methods of assessing renal function, such as serum creatinine and urea measurements, have limitations in the early detection of lesions^12^. The introduction of non-traditional biomarkers (BMs), such as Neutrophil Gelatinase-Associated Lipocalin (NGAL), Syndecan-1 (SYN-1), Angiotensin II (ANG-2), and Monocyte Chemoattractant Protein-1 (MCP-1), has shown potential for identifying renal and endothelial alterations in tropical diseases, enabling more sensitive and individualized clinical management. However, the behavior of these BMs in Chagas disease is not well understood^13,14,15,16^.

SYN-1 is a transmembrane proteoglycan (Type I) composed of heparan sulfate and belongs to the syndecan family of proteoglycans. Its elevated levels in plasma are generally associated with the loss of the endothelial glycocalyx and/or shedding of the renal epithelium and/or liver epithelium^17^. ANG-2 is a pro-inflammatory protein that can contribute to the pathogenesis of various inflammatory diseases, including cardiovascular diseases, especially congestive heart failure, pulmonary diseases, autoimmune diseases, and chronic kidney disease^18^. MCP-1, a crucial molecule, plays a significant role in modulating the activity of monocytes and macrophages and, being a chemokine, is responsible for attracting these cells to injured areas^16^. NGAL, a 25 kDa protein, is secreted by immune system cells and can be measured in plasma, urine, and other fluids. An important biomarker of acute kidney injury, NGAL is released after tubular damage and during renal regeneration, helping to predict the progression of renal failure and its complications. Additionally, NGAL is also useful in diagnosing cardiovascular diseases, showing high expression in injured cardiac tissue and atherosclerotic plaques, with serum concentrations reflecting the severity of heart failure and coronary artery disease^19–22^.

In the Brazilian Amazon, Chagas Disease (CD) presents distinct epidemiological characteristics compared to traditional vector-borne transmission areas. Notably, oral transmission that occurs through the consumption of food contaminated with *T. cruzi*, such as açaí, has emerged as the leading route of infection, frequently resulting in acute outbreaks in the region^6,23^. This epidemiological peculiarity contributes to the predominance of acute phase cases of the disease, characterized by clinical manifestations such as fever, headache, myalgia, facial edema, and cardiac dysfunction. The high prevalence of oral transmission together with a difficulty of accessing healthcare services, under diagnosis, and failure in clinical follow-up after treatment makes the Brazilian Amazon a region requiring special attention^23,24^. Based on data from *Sistema de Informação de Agravos de Notificação* (SINAN), approximately 3,800 cases were reported in this region between 2000 and 202, with the state of Pará accounting for 80% of the cases, highlighting the significant epidemiological impact of Chagas disease^25^.

In this region, although early detection of acute Chagas disease cases occurs through the malaria surveillance network^26,27^, clinical follow-up after treatment is often hindered by logistical challenges for most patients due to living in rural and remote areas with limited access to healthcare services ^23,24^. This limitation also impairs adequate monitoring of potential complications, such as renal lesions, which may significantly contribute to the morbidity and mortality associated with the disease. Furthermore, the impact of kidney impairment on patient prognosis remains an area scarcely explored within the context of Amazonian Chagas disease. Though known that renal injury can negatively influence outcomes, the incidence, and detailed characterization of such renal involvement in Chagas disease remains poorly documente^4^. In this context, our study aimed to investigate the expression and clinical relevance of non-traditional biomarkers of renal injury and endothelial activation—SYN-1, ANG-2, MCP-1, and NGAL— in patients with Chagas disease in the Brazilian Amazon region, with the goal of early detection of renal and vascular dysfunctions and improving clinical management of these patients.

## METHODS

### Study design, setting and patient selection

This was a cross-sectional study performed at Fundação de Medicina Tropical Doutor Heitor Vieira Dourado (FMT-HVD), a referral center for tropical and infectious diseases in the state of Amazonas, northern Brazil. The study population included patients with Chagas disease in the acute and chronic phases, native from the Brazilian Amazon.

Recruitment took place between July 2021 and July 2023 at the FMT-HVD Chagas Disease Outpatient Clinic. Here patients were grouped according to the stage of the disease. Group 1 – Acute phase: encompassing patients diagnosed through the malaria surveillance network in the Amazon, through thick blood smear exams, subdivided into: G1a (pre-treatment, at the time of diagnosis), G1b (post-treatment). Group 2 – Chronic phase: involving patients referred from cardiology clinics, blood donation and survey services, with diagnosis confirmed by serology, subdivided into: G2a (chronic indeterminate) and G2b (chronic cardiac).

Inclusion criteria was individuals with confirmed diagnosis of acute or chronic Chagas disease, of either sex, over 18 years of age and native to the Brazilian Legal Amazon were included. Group 1 - patients with positive parasitological diagnosis in acute phase, Group 2 - patients with reactive serology confirmed by at least two different diagnostic tests based on distinct methodologies.

Patients that were pregnant, living or traveling for more than 3 months to regions outside the Brazilian Legal Amazon, with a history of chronic kidney disease, organ transplantation, serious cardiovascular diseases (such as heart failure or thrombosis), or using non-steroidal anti-inflammatory drugs (NSAIDs) were excluded.

### Biological samples

Biological sample collection and laboratory analysis - Serum samples (4mL) were obtained in sterile tubes with separating gel ([BD Vacutainer® Serum Tubes](https://www.bd.com/en-us/products-and-solutions/products/product-page.367820)), urine (10mL) in specific bottles and whole blood (10mL) in tubes with EDTA (ethylenediaminetetraacetic acid) ([BD Vacutainer® Serum Tubes](https://www.bd.com/en-us/products-and-solutions/products/product-page.367820)). The biological samples were stored in aliquots at −20°C at Unidade de Entomologia Nelson Ferreira Fé (UENFF) at FMT-HVD, until use.

A complete blood count, biochemistry blood test (albumin, urea, creatinine, CPK, electrolytes), routine urine test and a 24-hour proteinuria were performed at the FMT-HVD Clinical Laboratory in the routine outpatient follow-up protocol. All patients underwent a kidney ultrasound, performed in a private clinic.

### Biomarker Assay

The quantification of biomarkers was performed using the ELISA (Enzyme-Linked Immunosorbent Assay) test, utilizing specific aliquots of serum and urine collected from patients at different time points. Commercial kits with specific antibodies for the target biomarkers were acquired: urinary NGAL (R & D Systems, ref: DY1757), urinary MCP-1 (R & D Systems, ref: DY279), angiopoietin-2 (R & D Systems, ref: DY923), and syndecan-1 (Abcam, ref: ab308538). The concentrations of urinary NGAL and MCP-1 were normalized by urinary creatinine from the same sample.

The methodology followed standardized protocols for the sandwich-type immunoenzymatic assay. Initially, 96-well plates were sensitized with a specific capture antibody then incubated for a predetermined time. After blocking to prevent nonspecific binding, urine and serum samples were added along with the standard curve for quantification. A detection antibody conjugated to an enzyme was then added, followed by the addition of the chromogenic substrate. The reaction was stopped by adding acid, and the absorbance was read in a spectrophotometer at 450 nm.

The experiments were conducted at the Pharmaceutical Bioprospecting and Clinical Biochemistry Laboratory (LBFBC) of the Federal University of Ceará.

### *T. cruzi* genetic analysis

Complementarily, *T. cruzi* DTUs were obtained from a database of the Chagas disease research group Dr. João Macias Frade. These strains were identified from blood samples from patients, using biomolecular techniques, according to the protocol used in the FMTHVD entomology laboratory^5^. DNA extraction from a peripheral blood sample (buffy coat) was performed following the PureLink™ Genomic DNA Mini kit protocol (Invitrogen, Life Technologies, California, USA).

### Evaluation of cardiac involvement

The assessment of alterations in the electrocardiogram followed the guidelines established by the Brazilian Society of Cardiology regarding the diagnosis and treatment of patients with Chronic Cardiopathy due to Chagas Disease (CCDC). Criteria used for cardiac alterations: a) ECG outside normal limits; b) 24-hour Holter — Ectopic activity (Sporadic — <200/24 hours; Mild — between 1 and 3% of the QRS number; Moderate — between 3% and 10% of the QRS number; Severe — between 10% and 30% of the QRS number; Very severe — above 30% of the QRS number); c) Echocardiogram: Reduced ejection fraction (FEVE) ≥50%; increased size of the right and left cardiac chambers; systolic dysfunction of the left ventricle.

### Ethical considerations

The Research Ethics Committee of the Fundação de Medicina Tropical Doutor Heitor Vieira Dourado approved this study under the CAAE number: 45886521.0.0000.0005, in accordance with Resolution No. 466/12 of the National Health Council of Brazil. Patients diagnosed with Chagas disease who participated in the study provided a signed written informed consent and completed a standardized questionnaire.

### Statistical analysis

Data analysis was performed using Stata v.14 (Stata Corp., USA). Variables were tested for normal distribution. Continuous parametric variables are reported as means ± SD and compared using analysis of variances (ANOVA). Continuous non-parametric variables are reported as medians and interquartile ranges and compared using Mann–Whitney U test with Tukey post-tests. Categorical data are reported as proportions and compared using Pearson’s qui-squared test. Spearman’s correlation (ρ) was applied between renal and endothelial biomarkers and proteinuria. A p-value <0.05 was considered statistically significant for all comparisons.

## RESULTS

### Socio-epidemiological and clinical laboratory aspects

The study included 78 patients, with a mean age of 43.5 ± 16.4 years with 43(55.1%) being male patients. The majority, 73(93.6%), came from 14 municipalities in the state of Amazonas; and 58(74.4%) were infected with *T. cruzi* during outbreaks due to oral transmission. Patient stratification by groups and disease duration, revealed 67(85.9%) patients were in G1, most of whom, 53 (67.9%), were in G1b (post-treatment). The average clinical follow-up time was 5.9 ± 5.2 years after diagnosis, ranging from 6 months to 19 years. In G2, 11 patients were evaluated, of which G2b (chronic cardiac phase) was the largest subgroup with 7 (63.6%) patients. From the G1 category, *T. cruzi* lineage identification was performed in only 56(83.6%) patients, of whom the majority, 54/56 (96.4%), had TcIV. Regarding comorbidities, these were observed in 20 (25.6%) patients, with systemic arterial hypertension (SAH) being the most prevalent (17; 85.0%). Most of the comorbities (14; 70.0%) were among individuals in the G1b category (Table 1).

**Table 1.** Socio-epidemiological and clinical characteristics of patients with Chagas disease.

| Variables | Total<br>(n=78)<sup>1</sup> | G1a<br>(n=14)<sup>1</sup> | G1b<br>(n=53)<sup>1</sup> | G2a<br>(n=4)<sup>1</sup> | G2b<br>(n=7)<sup>1</sup> | P |
| --- | --- | --- | --- | --- | --- | --- |
| Age, mean (SD), years | 43.5 ± 16.4 | 40.2 ± 14.7 | 42.8 ± 17.3 | 49.5 ± 8.4 | 51.9 ± 15.2 | 0.511<sup>2</sup> |
| Sex |  |  |  |  |  |  |
| Male | 43 (55.1) | 12 (85.7) | 25 (47.2) | 3 (75.0) | 3 (42.9) | 0.052<sup>3</sup> |
| Female | 35 (44.9) | 2 (14.3) | 28 (52.8) | 1 (25.0) | 4 (57.1) |  |
| Origin | | | | | | $<0.001^3$ |
| Amazonas State – 14 municipalities | | | | | | $<0.001^3$ |
| Amaturá-AM | 9 (11.5) | 9 (64.3) | - | - | - |  |
| Barreirinha-AM | 15 (15.3) | 1 (7.1) | 14 (26.4) | - | - |  |
| Carauari-AM | 13 (13.3) | - | 12 (22.6) | 1 (25.0) | - |  |
| Careiro-AM | 1 (1.0) | - | - | - | 1 (14.3) |  |
| Coari-AM | 10 (10.2) | - | 10 (18.9) | - | - |  |
| Eirunepé-AM | 1 (1.0) | - | 1 (1.9) | - | - |  |
| Ipixuna-AM | 3 (3.1) | 3 (21.4) | - | - | - |  |
| Lábrea-AM | 4 (4.1) | - | 4 (7.6) | - | - |  |
| Manaus-AM | 28 (28.6) | - | 4 (7.6) | 3 (75.0) | 1 (14.3) |  |
| Rio Preto da Eva-AM | 1 (1.0) | 1 (7.1) | - | - | - |  |
| Santa Isabel R Negro-AM | 2 (2.0) | - | 2 (3.8) | - | - |  |
| Tabatinga-AM | 1 (1.0) | - | 1 (1.9) | - | - |  |
| Tefé-AM | 2 (2.0) | - | 1 (1.9) | - | 1 (14.3) |  |
| Uarini-AM | 3 (3.1) | - | 3 (5.7) | - | - |  |
| Pará – 3 municipalities |  |  |  |  |  | 0.223<sup>3</sup> |
| Belém-PA | 1 (1.0) | - | - | - | 1 (14.3) |  |
| Juruti-PA | 1 (1.0) | - | - | - | 1 (14.3) |  |
| Marajó-PA | 1 (1.0) | - | 1 (1.9) | - | - |  |
| Maranhão – 2 municipalities |  |  |  |  |  | na |
| Axixá-MA | 1 (1.0) | - | - | - | 1 (14.3) |  |
| Pedreiras – MA | 1 (1.0) | - | - | - | 1 (14.3) |  |
| **Case Type** |  |  |  |  |  | < 0.001<sup>3</sup> |
| Chronic | 11 (11.2) | - | - | 4 (100) | 7 (100) |  |
| Isolated | 9 (9.2) | 2 (14.3) | 7 (13.2) | - | - |  |
| Outbreak | 58 (59.2) | 12 (85.7) | 46 (86.8) | - | - |  |
| **Follow-up time (years)** | 5.9 ± 5.2 | - | 7.4 ± 4.9 | 6.5 ± 2.1 | 6.6 ± 5.2 | 0.282<sup>3</sup> |
| **DTU *T. cruzi*** | **n = 56 (71.8)** | **N = 13 (92.8)** | **n = 43 (81.1)** | **n = 0** | **n = 0** | 0.731<sup>3</sup> |
| TcI | 1 (1.8) | - | 1 (2.3) | - | - |  |
| TcIV | 54 (96.4) | 13 (100) | 41 (95.4) | - | - |  |
| Z3 (TcIII/TcIV) | 1 (1.8) | - | 1 (2.3) | - | - |  |
| **Comorbidities** | **n = 20 (25.6)** | **n = 2 (14.3)** | **n = 14 (26.4)** | **n = 0** | **n = 4 (57.1)** | 0.129<sup>3</sup> |
| Hypertension | 13 (65.0) | 1 (50.0) | 8 (57.2) | - | 4 (100.0) |  |
| Diabetes mellitus | 1 (5.0) | 1 (50.0) | - | - | - |  |
| SAH/DM | 4 (20.0) | - | 4 (28.6) | - | - |  |
| Asthma | 1 (5.0) | - | 1 (7.1) | - | - |  |
| Shehann Syndrome | 1 (5.0) | - | 1 (7.1) | - | - |  |

\*DM – Diabetes mellitus. SAH – Systemic arterial hypertension. <sup>1</sup>n (%); <sup>2</sup>ANOVA; <sup>3</sup>Pearson chi-square test

### Cardiac alterations

All patients underwent an ECG, and 25 (32%) showed some type of abnormality, including bundle branch blocks and ST/T segment changes; 48 (61.5%) had an echocardiogram, and 5 (10%) exhibited abnormalities; 51 (65.4%) were submitted to 24-hour Holter monitoring, and 6 (11.8%) showed abnormalities (Table 2).

**Table 2.**
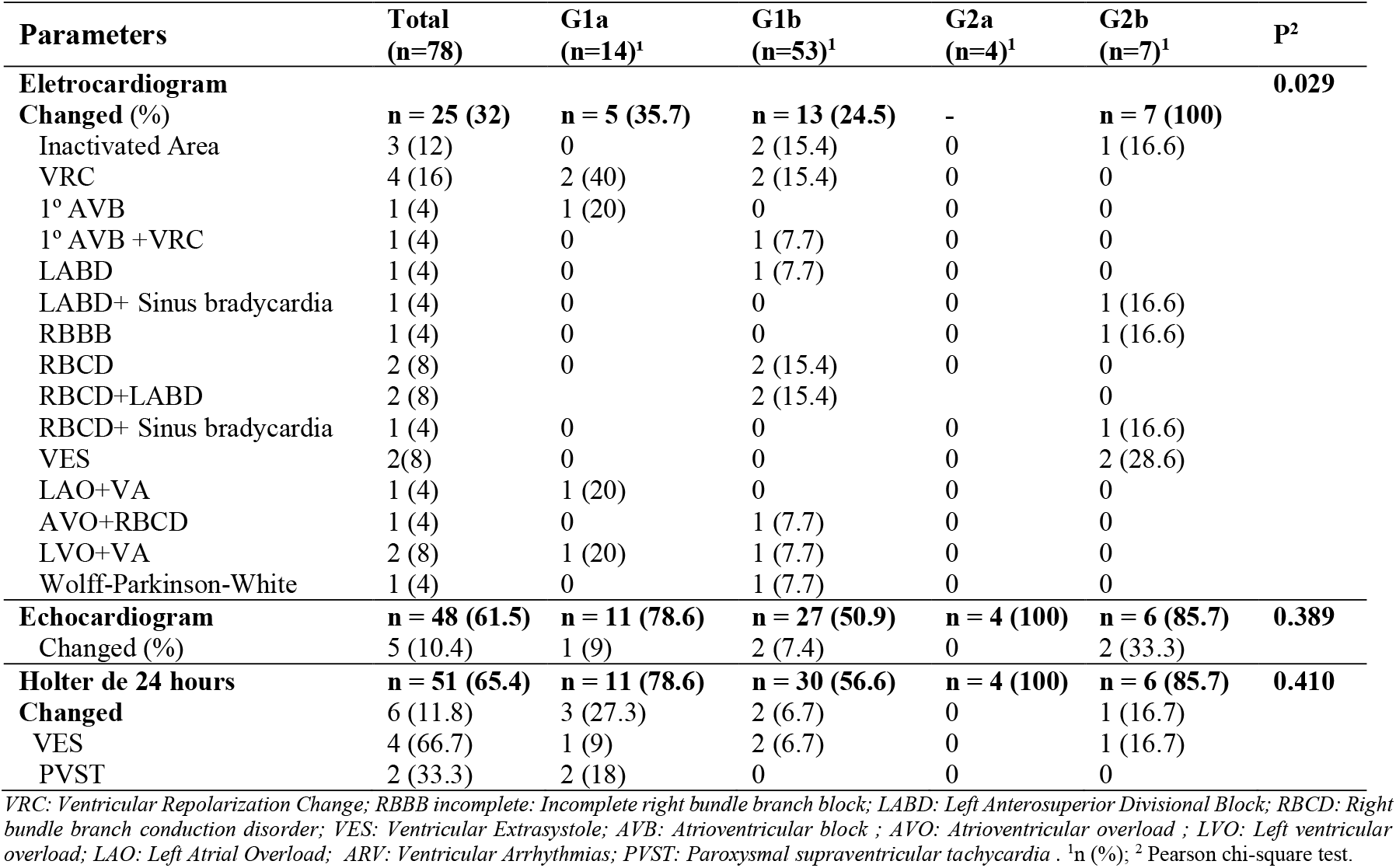
Presence of cardiac alterations in patients with Chagas disease in the Amazon.

### Routine additional examinations

From the routine laboratory tests performed, no patient presented anemia. However, there was a statistically significant difference in platelet count, leukocytes, glucose, glycated hemoglobin (HbA1c), high-density lipoprotein cholesterol (HDL), aspartate aminotransferase (AST), alanine aminotransferase (ALT), gamma-glutamyl transferase (GGT), and creatine kinase (CK) (all p<0.05) Blood glucose and HbA1c levels were elevated in group G1b, with averages of 115 ± 66.3 and 6.3 ± 2.3, respectively. In the type 1 urine analysis, samples were collected from 44 (56.4%) patients, of whom 5 (11.4%) exhibited microscopic hematuria with more than 3 red blood cells per field. In these cases, an analysis was conducted to detect erythrocyte dysmorphisms in isolated urine samples, all of which yielded negative results. Neither proteinuria nor cylindruria was observed in any of the patients (Table 3). A renal ultrasound imaging exam was performed in 39 (50%) patients, and none showed structural abnormalities.

**Table 3.** Profile of routine laboratory tests of patients with Chagas disease.

| Variables | Total<br>(n=78)<sup>1</sup> | G1a<br>(n=14)<sup>1</sup> | G1b<br>(n=53)<sup>1</sup> | G2a<br>(n=4)<sup>1</sup> | G2b<br>(n=7)<sup>1</sup> | P |
| --- | --- | --- | --- | --- | --- | --- |
| **Hemogram** |  |  |  |  |  |  |
| Hemoglobin (g/dL) | 14.3 ± 1.7 | 13.2 ± 1.4 | 14.4 ± 1.6 | 16.4 ± 0.3 | 15 ± 1.2 | 0.534<sup>2</sup> |
| Hematocrit (%) | 43.4 ± 5.0 | 39.9 ± 4.2 | 43.6 ± 4.3 | 49.1 ± 0.2 | 46.8 ± 6.4 | 0.107<sup>2</sup> |
| Platelets (/10<sup>3</sup>) | 261 ± 11 | 323 ± 19 | 252 ± 59 | 225 ± 53 | 208 ± 64 | < 0.001<sup>2</sup> |
| Leucocytes (/10<sup>3</sup>) | 7.4 ± 2.5 | 7.9 ± 2.3 | 7.6 ± 2.4 | 4.6 ± 5.8 | 6.2 ± 8 | 0.048<sup>2</sup> |
| **Biochemistry tests** |  |  |  |  |  |  |
| Glucose (g/dL) | 108.9 ± 52.6 | 101.7 ± 15.6 | 115 ± 66.3 | 96 ± 2.6 | 97.6 ± 6.2 | < 0.001<sup>2</sup> |
| HbA1c (%) | 6.1 ± 2.0 | - | 6.3 ± 2.3 | 5.6 ± 0.2 | 5.4 ± 0.1 | < 0.001<sup>2</sup> |
| Cholesterol total (mg/dL) | 192.1 ± 42.4 | 162.9 ± 21.5 | 200 ± 47.2 | 188 ± 26.6 | 198.7 ± 31.8 | 0.076<sup>2</sup> |
| LDL-cholesterol (mg/dL) | 122.5 ± 37 | 97.7 ± 23.5 | 131 ± 40.6 | 117 ± 14 | 123.5 ± 28.1 | 0.137<sup>2</sup> |
| HDL-cholesterol (mg/dL) | 42.3 ± 22.3 | 28.9 ± 7.6 | 41.7 ± 9.8 | 51 ± 29.3 | 62.8 ± 52.6 | < 0.001<sup>2</sup> |
| Triglycerides (mg/dL) | 156.8 ± 86.1 | 182 ± 105.2 | 146 ± 84.6 | 127 ± 53.4 | 178.8 ± 74.3 | 0.666<sup>2</sup> |
| AST (U/L) | 35.6 ± 33.0 | 63.5 ± 52.3 | 28.6 ± 21.5 | 26.3 ± 3.1 | 21.2 ± 6.4 | < 0.001<sup>2</sup> |
| ALT (U/L) | 45.6 ± 47.5 | 85 ± 85.6 | 35.3 ± 18.5 | 33.3 ± 11.6 | 28.4 ± 24.8 | < 0.001<sup>2</sup> |
| Alkaline phosphatase (U/L) | 258 ± 106.4 | 360 ± 131.3 | 227 ± 62.8 | 227.8 ± 62.8 | 203.3 ± 63.5 | 0.038<sup>2</sup> |
| Gamma GT (U/L) | 55.6 ± 52.7 | 108.2 ± 78.1 | 37.1 ± 16.3 | 27.5 ± 14.8 | 35.3 ± 26.3 | < 0.001<sup>2</sup> |
| Albumin (g/dL) | 4.4 ± 0.3 | 4.1 ± 0.3 | 4.6 ± 0.2 | 4.3 ± 0.4 | 4.5 ± 0.5 | 0.469<sup>2</sup> |
| Creatine kinase (U/L) | 143 ± 321.1 | 46.8 ± 13.4 | 96.8 ± 40.6 | - | 552 ± 820.6 | < 0.001<sup>2</sup> |
| Potassium (mEq/L) | 4.4 ± 0.4 | 4.4 ± 0.5 | 4.4 ± 0.4 | 4.4 ± 0.9 | 4.3 ± 0.3 | 0.135<sup>2</sup> |
| Sodium (mEq/L) | 139 ± 3.4 | 138.3 ± 3.7 | 139.2 ± 3.6 | 140.7 ± 2.5 | 138.3 ± 2.7 | 0.822<sup>2</sup> |
| Calcium (mg/dL) | 10.1 ± 0.6 | 9.8 ± 1.2 | 10.3 ± 0.4 | - | 9.9 ± 0.6 | 0.395<sup>2</sup> |
| **Urinalysis, n (%)** | **44 (56.4)** | **10 (71.4)** | **26 (49.1)** | **2 (50)** | **6 (85.7)** |  |
| Density | 1016 ± 8.0 | 1017 ± 7.5 | 1017 ± 8.1 | 1015 ± 14.1 | 1013 ± 8.2 | 0.830<sup>2</sup> |
| pH | 6.1 ± 0.3 | 5.9 ± 0.3 | 6.1 ± 0.2 | 6.5 ± 0.7 | 6.1 ± 0.2 | 0.130<sup>2</sup> |
| Hemoglobin | 9 (20.5) | - | 7 (26.9) | - | 2 (33.3) | 0.225<sup>3</sup> |
| Glucose | 3 (6.8) | - | 2 (7.7) | - | 1 (16.7) | 0.609<sup>3</sup> |
| Pyocytes | 17 (38.6) | 4 (40) | 13 (50) | - | - | 0.091<sup>3</sup> |
| Red Blood Cells | 5 (11.4) | - | 4 (15.4) | - | 1 (16.7) | 0.547<sup>3</sup> |
| Crystals | 10 (22.7) | 4 (40) | 6 (23.1) | - | - | 0.256<sup>3</sup> |
| Bacteria | 19 (44.2) | 5 (50) | 14 (56) | - | - | 0.048<sup>3</sup> |

AST – Aspartate aminotransferase. ALT – Alanine aminotransferase. Gamma GT - Gamma glutamyltransferase. HbA1c – Glycated hemoglobin. <sup>1</sup>n (%);
<sup>2</sup>ANOVA; <sup>3</sup> Pearson chi-square test

### Renal and endothelial biomarkers

The estimated glomerular filtration rate (eGFR) calculated using serum creatinine (ml/min/1.73 m^2^), based on the CKD-EPI equation (Chronic Kidney Disease Epidemiology Collaboration), remained within normal values in all analyzed groups, with a mean of 109.4 ± 10.2. It was observed that 7 out of 78 (9.0%) patients exhibited hyperfiltration with an eGFR > 120 mL/min/1.73 m^2^, of whom 2 had comorbidities (hypertension/diabetes mellitus) and one had Chagas’ cardiomyopathy. The 24-hour proteinuria samples, in turn, were within the considered normal range (Table 4).

**Table 4.** Profile of renal and endothelial biomarkers of patients.

| Variables | Control<br>(n=20)<sup>1</sup> | G1a<br>(n=14)<sup>1</sup> | G1b<br>(n=53)<sup>1</sup> | G2a<br>(n=4)<sup>1</sup> | G2b<br>(n=7)<sup>1</sup> | P |
| --- | --- | --- | --- | --- | --- | --- |
| **Biomarkers of kidney function** |  |  |  |  |  |  |
| eGFR >120 | 0 (NA%) | 2 (14.3) | 3 (5.7) | 1 (25) | 1 (14.3) | 0.441<sup>2</sup> |
| Creatinine (mg/dL) | NA ± NA | 0.9 ± 0.1 | 0.8 ± 0.2 | 1.0 ± 0.1 | 0.9 ± 0.2 | 0.577<sup>3</sup> |
| Urea | NA ± NA | 32.8 ± 6.9 | 32.8 ± 10.8 | 29.7 ± 7.2 | 33.3 ± 7.8 | 0.953<sup>3</sup> |
| eGFR-Cr<br>(mL/min/1.73m<sup>2</sup>) | NA ± NA | 109.4 ± 10.2 | 101.9 ± 19.6 | 95.3 ± 15.3 | 95.2 ± 12.7 | 0.342<sup>3</sup> |
| 24h protein (g/24h) | NA ± NA | 93.1 ± 56 | 55.6 ± 31.3 | 72.4 ± 44.2 | 68.8 ± 20.6 | 0.170<sup>3</sup> |
| **Biomarkers of kidney injury** |  |  |  |  |  |  |
| NGAL corrected,<br>(pg/mL-Cr) | 8.5<br>[4.8 - 23.1] | 22.0<br>[8.5 - 27.7] | 21.2<br>[10.6 - 41.9] | 33.3<br>[11.3 - 58.1] | 28.1<br>[8.5 - 37.7] | 0.064<sup>4</sup> |
| MCP-1 corrected,<br>(ng/mg-Cr) | 52.8<br>[34.0 - 77.6] | 266.8<br>[153.4 - 440.1] | 176.3<br>[76.3 - 254.1] | 70.4<br>[31.7 - 113.2] | 55.7<br>[30.2 - 97.1] | <0.001<sup>4</sup> |
| **Biomarkers of endothelial injury** |  |  |  |  |  |  |
| Syn-1<br>(ng/mL) | 42.9<br>[31.3 - 56.9] | 402.3<br>[211.7 - 6988.0] | 67.3<br>[34.7 - 2892.2] | 2682.7<br>[1083.4 - 3289.1] | 179.1<br>[49.3 - 2600.0] | 0.001<sup>4</sup> |
| Angiopoietin-2<br>(ng/mL) | 1.8<br>[0.9 - 3.1] | 0.1<br>[0.0 - 2.7] | 1.0<br>[0.7 - 1.4] | 1.2<br>[0.7 - 4.0] | 1.4<br>[1.2 - 2.2] | 0.156<sup>4</sup> |

eGFR – Estimated glomerular filtration rate. eGFR-Cr – Estimated glomerular filtration rate for creatinine. NGAL – Neutrophil Gelatinase-Associated Lipocalin; MCP-1 – Monocyte chemoattractant protein-1, Syn-1 – Syndecan-1. <sup>1</sup>n(%); <sup>2</sup>Pearson's chi-square test; <sup>3</sup>ANOVA test; <sup>4</sup>Kruskal-Wallis rank sum test.

Regarding the renal biomarkers, NGAL was higher in the G2a group, with 33.3 [11.3 - 58.1] ng/mg-Cr, but without statistical significance (p=0.064). The inflammatory marker MCP-1 showed increased urinary levels in group G1a, with 266.8 [153.4 - 440.1] ng/mg-Cr, compared to the other groups, with a statistically significant difference (p<0.001). Concerning the endothelial biomarkers, SYN-1 exhibited increased expression in group G2a, ranging from 1,083.4 to 3,289.1 ng/mL. However, in the pre-treatment acute group G1a, it showed wide variability, oscillating from 211.7 to 6,988.0 ng/mL. When stratifying patients by groups, it was observed that SYN-1 levels were elevated in subgroups G1a (pre-treatment) and G2a (indeterminate), presenting a statistically significant difference (p=0.001). The vascular biomarker ANG-2 showed a slight increase in patients from group G2, with an average of 1.2 (7-4.0) in G2a and an average of 1.4 (1.2-2.2) in G2b, when compared to G1 patients, without a statistically significant difference (Table 4).

### Comorbidities

Comparing patients with comorbidities (n=20) to those without comorbidities (n=58), Syn-1 levels were significantly higher in patients with comorbidities, with an approximate ratio of 31:1. Contrastingly, ANG-2 remained within normal range among those without comorbidities but slightly elevated in those with comorbidities, suggesting a potential association between ANG-2 expression and underlying pathological conditions. Analysis of ANG-2 levels revealed that patients with hypertension (SAH) or combined SAH and diabetes mellitus exhibited elevated concentrations, with median values of [1.0–2.4] ng/mL and 1.5 [1.3–3.5] ng/mL, respectively. In contrast, patients without comorbidities had a lower median ANG-2 level of 0.9 [0.5–1.5] ng/mL. Urinary NGAL adjusted for creatinine showed higher concentration in patients without comorbidities compared to those with comorbidities, although the difference did not reach statistical significance (p=0.871). Meanwhile, urinary MCP-1 remained similar between the two groups (0.15 ng/mg-Cr – comorbidities v/s 0.14 ng/mg-Cr – no comorbidities, p=0.894) (Table 5).

**Table 5.**
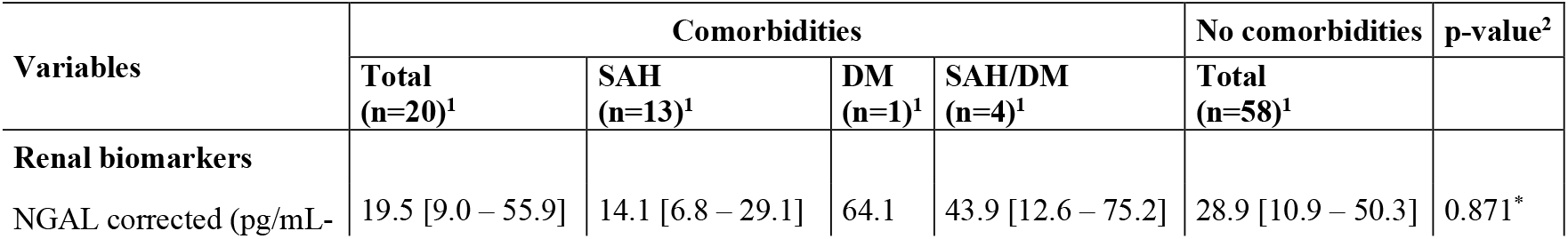
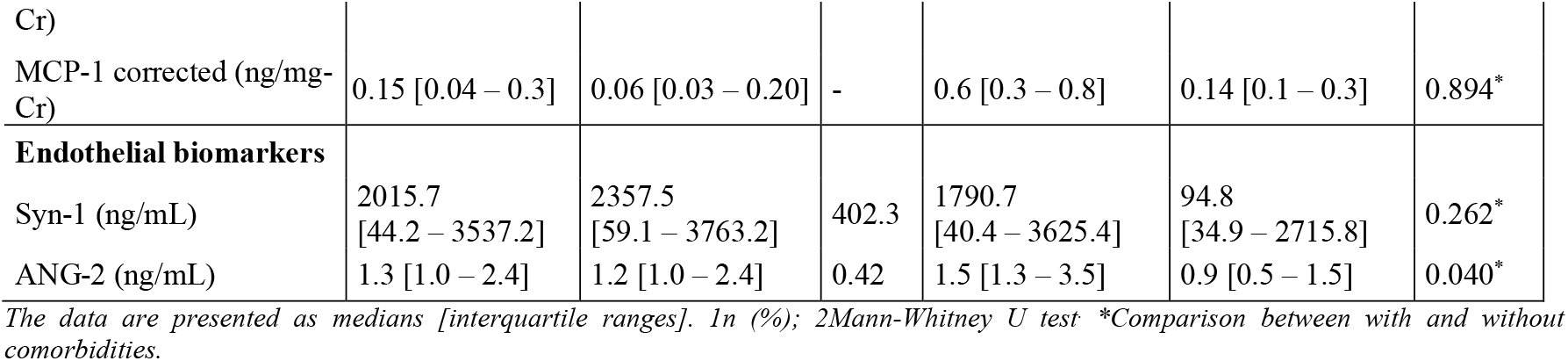
Profile of biomarkers of kidney and endothelial injury in the presence of comorbidities.

### Relationship between the new biomarkers and traditional renal markers

Traditional renal function markers - including creatinine, urea, 24-hour proteinuria and GFR-were analyzed alongside serum levels of SYN-1, ANG-2, MCP-1, and urinary NGAL. No significant correlations were observed between creatinine or urea and any of the measured biomarkers (Figures 1 and 2). Although 24-hour proteinuria remained within normal ranges, a weak, non-significant positive correlation was noted between MCP-1 and urinary NGAL (Figure 3). GFR showed a very weak negative correlation with urinary NGAL, but no correlation with SYN-1, ANG-2, or MCP-1 (Figure 4). These findings suggest potential subclinical alterations in renal or endothelial function that are not detectable by conventional renal markers.

**Figure 1.**
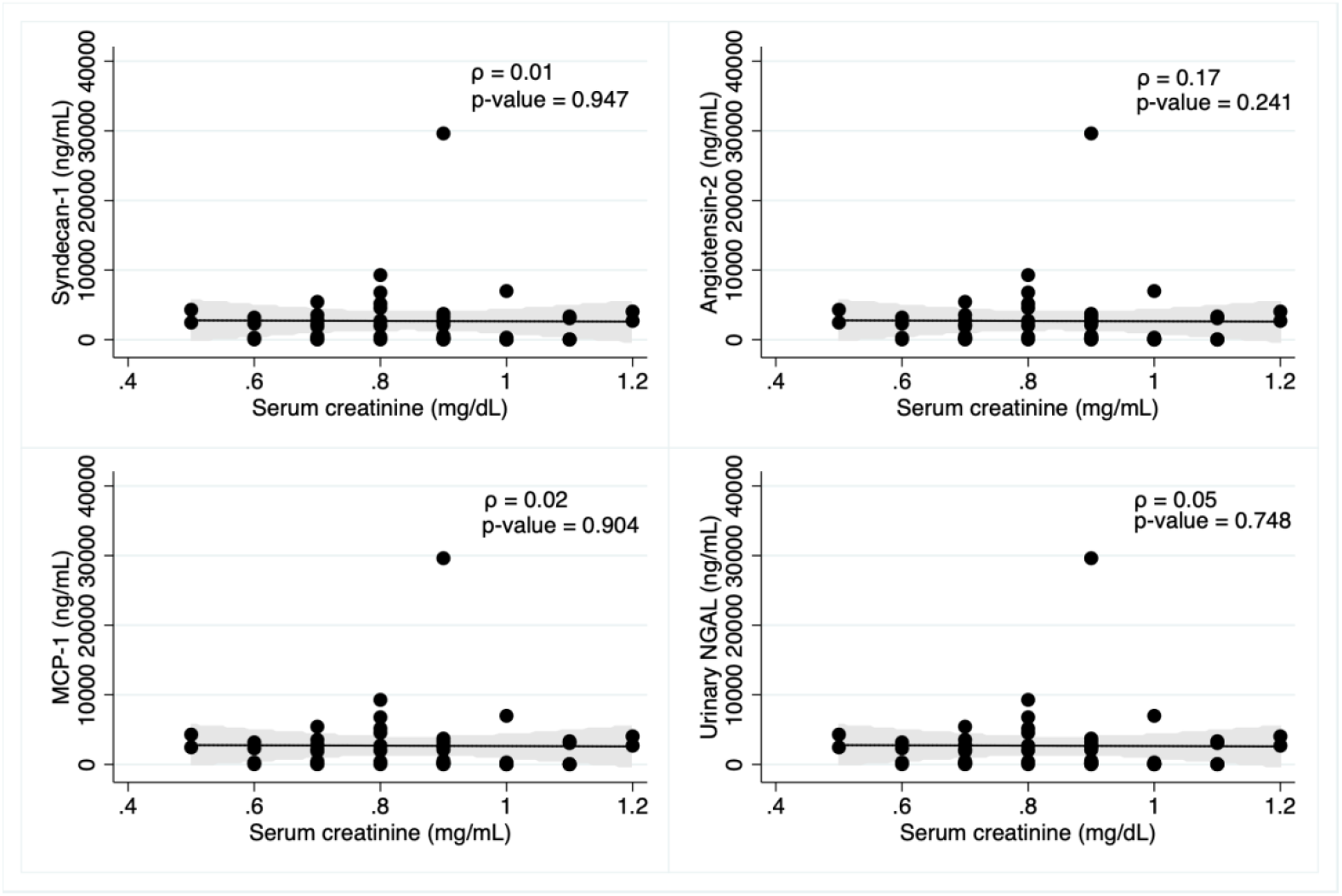
Pearson’s correlation (ρ) between biomarker levels and serum creatinine levels.

**Figure 2.**
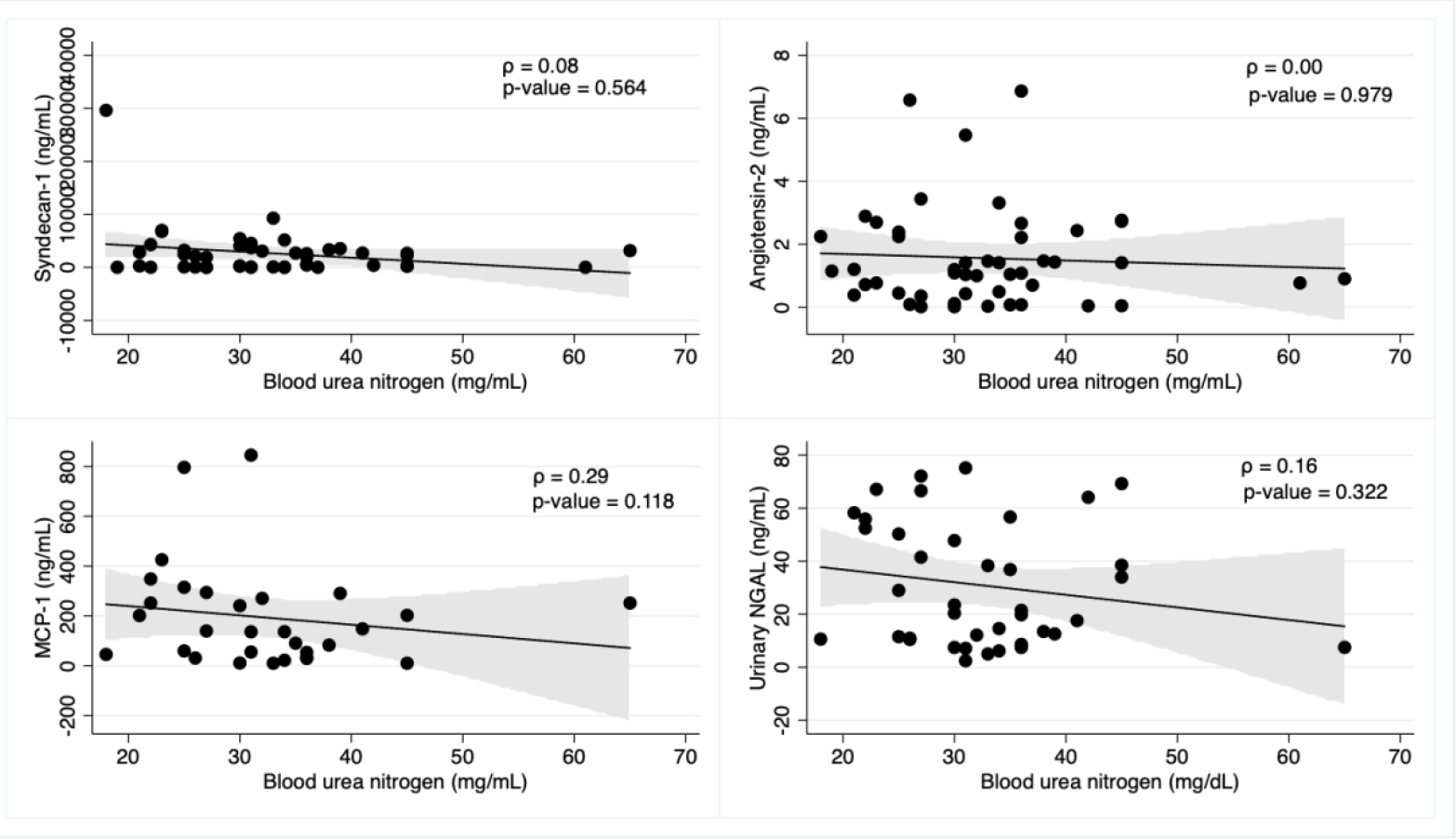
Spearman’s correlation (ρ) between biomarker levels and urea levels.

**Figure 3.**
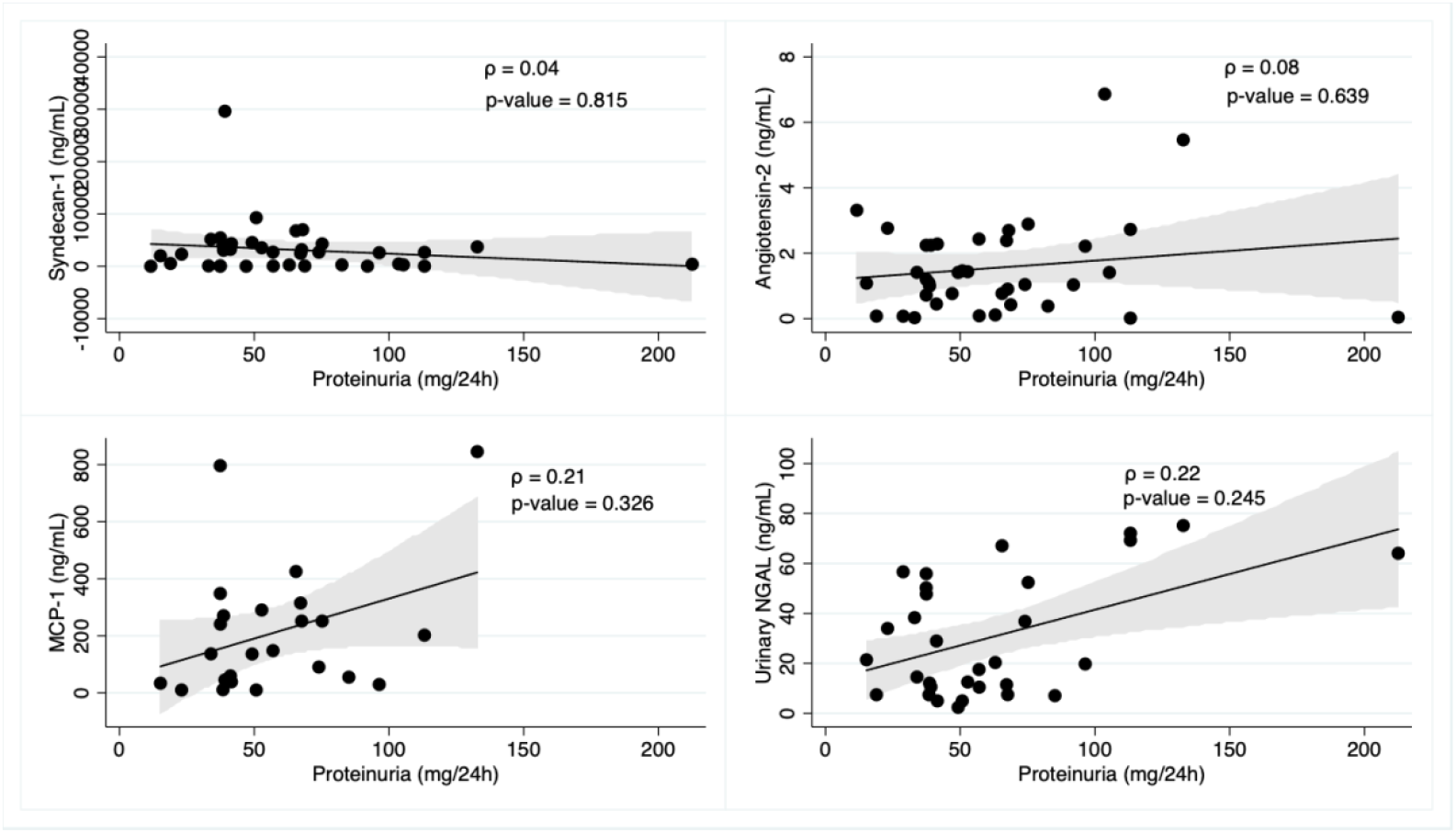
Spearman’s correlation (ρ) between biomarker levels and proteinuria.

**Figure 4.**
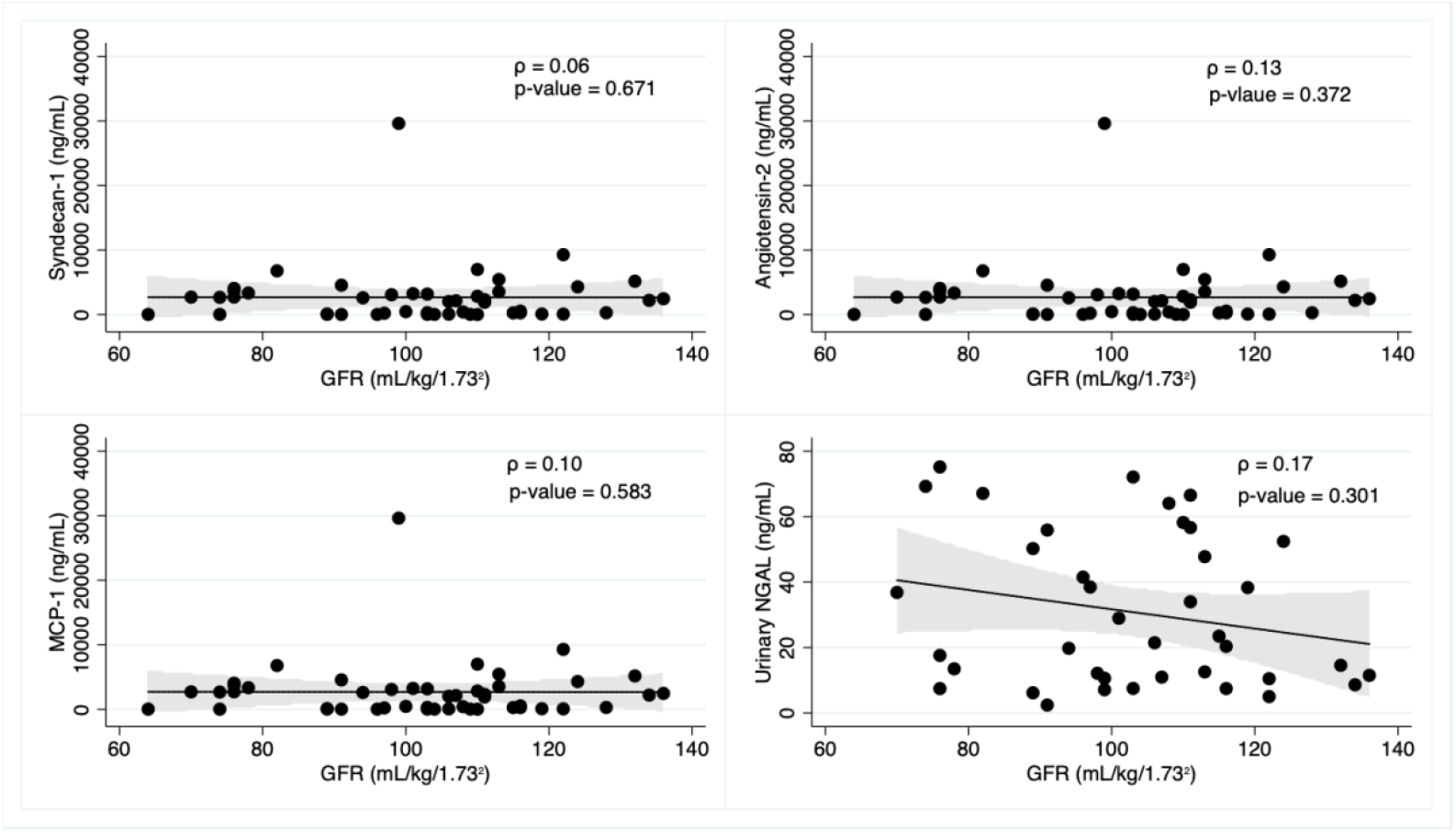
Spearman’s correlation (ρ) between biomarker levels and glomerular filtration rate.

## DISCUSSION

This study highlights the particularities of Chagas disease (CD) in the Amazon, a region characterized by vast geographic distances, limited access to health services, and other socioeconomic barriers that hinder the adequate monitoring of patients with Chagas disease. In this area, oral transmission prevails ^28,29,30^ especially associated with the consumption of contaminated foods, such as *açaí* ^31^, which differentiates the local epidemiological profile from traditional endemic regions where vector transmission is predominant. The increased severity of acute Chagas disease cases is believed to be associated with a higher *T. cruzi* inoculum via the oral transmission compared to vectorial transmission^32^. This route of infection may influence both clinical outcomes and immune responses, potentially affecting biomarker expression and the overall progression of the disease.

In Amazonas state, a pioneering study evaluated patients by means of cardiovascular magnetic resonance imaging, with an average follow-up of 5.2 years after diagnosis and treatment of the disease, identifying myocardial injury in 18% of cases ^33^. This follow-up period allowed observation of both clinical progression and cardiac alterations, highlighting the importance of continuous monitoring to understand the long-term impacts of the infection and involvement of other organs. However, this type of follow-up is especially challenging in the Amazon, as few patients return to health services for reassessment of their condition. Notably, although nearly half of the participants had been living with the disease for over 10 years, none had received any clinical follow-up after their initial treatment.

In this study, the participants’ average age of 40 years was within a range commonly reported in CD studies^33–36^. This age-group corresponds to an economically active population being subsequently affected resulting in significant socioeconomic impacts due to their impairment from the disease. The low frequency of patients in the chronic phase and with severe cardiac involvement recorded in the region, in this context, may indicate a milder profile of the CD. However, the difficulty in doing a longitudinal monitoring of patients, evidenced by the absence of follow-up for clinical evaluations in more than 40% of cases, compromises the epidemiological and clinical assessment of disease progression.

Due to the systemic nature of CD, especially in the acute phase, the most severe clinical manifestations include cardiac and neurological involvement that increase the risk of morbidity and mortality ^23,34,35^. In the acute phase, clinical symptoms primarily include fever, headache, myalgia, facial and lower limb edema, abdominal pain, and myocarditis ^23^. These clinical manifestations can be severe and differ in certain aspects when compared to the classic vector-borne transmission manner ^6^. The low morbidity and mortality commonly observed in patients from Amazonas may be related to early detection and more timely treatment; however, clinical follow-up of patients remains a major challenge due to the low rate of patient return, limited access to healthcare services, and low adherence to post-treatment monitoring ^23,36^. Cardiac alterations are also detectable in the post-treatment period ^34^, and patients diagnosed in the chronic cardiac phase are fewer and exhibit less severity compared to those from other traditionally endemic areas ^36^.

The present study included patients from all reported acute Chagas disease outbreaks in Amazonas state between 2004 and 2022, along with isolated cases of suspected vectorial transmission and chronic CD. Many participants resided in remote, hard-to-reach areas, and 23 were only detected through active outreach in their home municipalities. A key concern is that none of these patients had received clinical follow-up after treatment, primarily due to geographical isolation of their communities.

Considering that infection by *T. cruzi* is associated with inflammatory responses affecting the cardiovascular system, both in acute and chronic cases, and that additional factors such as renal involvement may also occur, it is important to remember that this inflammatory process can also trigger kidney disease^11^. In Amazonas a patient with acute Chagas disease, on the fifth day of evolution, presented cardiorenal syndrome characterized by myocarditis, a reduced ejection fraction (45%), and renal impairment evidenced by creatinine of 1.8 mg/dl, urea of 69 mg/dl, and GFR of 30 ml/min/1.73m^2 37^. These observations influenced the need to conduct the present study. In this context, we present novel findings suggesting potential renal alterations in patients with Chagas disease, based on the evaluation of early biomarkers of kidney injury and endothelial dysfunction, including NGAL, MCP-1, ANG-2, and SYN-1. Here, we demonstrate their potential value in detecting subtle functional changes already happening in the initial stages of the disease, anticipating alterations identifiable by traditional methods.

Experimental studies have documented biochemical and histological alterations during the acute phase of Chagas disease, including elevated blood urea, proteinuria, kidney injury molecule-1 (KIM-1), tubular necrosis, hypercellularity, mesangial congestion, vasculitis, and fibrosis^3^. In the present study, despite traditional markers of renal function, such as creatinine, urea, proteinuria, and GFR, being within normal parameters, an elevation in renal and endothelial biomarkers suggested subclinical renal and endothelial impairment in the studied population. This indicates that, even without evident clinical alterations in renal function, there are biochemical and molecular signs of renal injury and endothelial dysfunction associated with *T. cruzi* infection. Elevated levels of renal and endothelial biomarkers, such as NGAL, MCP-1, SYN-1 and ANG-2, observed in both acute Chagas disease (ACD) and chronic Chagas cardiomyopathy (CCC) patients underscores their value in early detection of renal complications in CD, especially in regions like the Amazon, where long-term clinical follow-up of patients may be challenging.

The presence of comorbidities with CD are capable of intensifying renal and cardiovascular impairment as demonstrated by the elevated levels of endothelial biomarkers observed. This further reinforces the need for an integrated health approach that addresses not only *T. cruzi* infection but also comorbid conditions that may impact the clinical progression and prognosis of affected individuals. The significant elevation of SYN-1 in patients with comorbidities suggests a loss of endothelial glycocalyx integrity, reflecting continuous or progressive endothelial damage that may contribute to disease progression and impairment of renal microcirculation. This marker has been associated with other infectious diseases involving the kidneys, such as leptospirosis, supporting the hypothesis that inflammation and silent endothelial damage constitute common and potentially silent pathogenic mechanisms^38^.

The pathophysiology of renal injury in Chagas disease involves an intricate interaction between immunological, autoimmune, and inflammatory processes, including glomerulonephritis and systemic manifestations resulting from the parasite infection, as well as direct damage to renal tissue. *T. cruzi* has the ability to infect various cells, including renal cells, leading to functional and structural alterations in the kidneys under different conditions. This consequently triggers an inflammatory and immune response that stimulates the production of pro-inflammatory mediators, causing direct cellular injury and systemic activation of the renin-angiotensin-aldosterone system^4,5,9,10,39,40^. These mechanisms can culminate in tubular injury, glomerulonephritis, and fibrosis, as demonstrated in experimental models ^3,11,41,42,43,44,45^. Understanding such processes and mechanisms is of utmost importance in the Amazon region, more so because of acute CD predominance and difficulties in accessing medical care hinder early diagnosis and management of the disease. The complexity of these pathophysiological mechanisms underscores the need for careful monitoring and integrated approaches to identify early possible renal damage among CD patients.

Urinary SYN-1 increases as acute kidney injury progresses highlighting its diagnostic and prognostic value. Understanding the relationship between SYN-1 expression and renal function can provide important insights into the underlying mechanisms of kidney injury and potential therapeutic targets. Similarly, this biomarker is associated with cardiovascular functions^14^. There are indications that lower plasmic levels of SYN-1 are associated with cardiovascular events and linked to overall mortality in patients undergoing hemodialysis^17^. Therefore, SYN-1 has the potential of an early biomarker of acute kidney injury to assess the severity of renal damage, in addition to cardiac functions and other diseases. However, this is the first clinical evaluation study using these markers in Chagas disease across its different clinical stages. Another biomarker of interest is ANG-2, an endothelial-derived protein that plays an important role in regulating vascularization and inflammation. ANG-2 biomarkers have been studied particularly for the correlations between chronic kidney disease and cardiovascular events^18^.

A protein associated with cardiovascular diseases, ANG-2 can be leveraged as an important indicator of chronic kidney disease (CKD) which predisposes patients to increased risk of cardiovascular diseases. Imbalance in ANG-2 levels has adverse effects on cardiovascular function, causing vascular dysfunction, inflammation, and remodeling. This directly influences the health of blood vessels, the heart, and the kidneys, as well as exerting indirect control over immune responses. A study investigating the association between ANG-2 and subclinical cardiovascular alterations in patients with CKD demonstrated that elevated ANG-2 levels correlated with increased left ventricular mass and wall thickness. Moreover, patients with high ANG-2 concentrations exhibited a higher risk of cardiovascular events and mortality, supporting the relevance of ANG-2 in the comorbidities observed in this population. In individuals with advanced CKD (stages 3b–5), elevated ANG-2 was also linked to an increased likelihood of initiating renal replacement therapy (RRT). Elucidating the mechanisms driving this association may inform the development of targeted interventions aimed at slowing CKD progression and delaying the need for RRT^15^.

MCP-1, an inflammatory urinary chemokine, was elevated in the acute pre-treatment phase, highlighting its role as an early marker of kidney injury and local inflammation, even in the absence of changes in traditional markers. Its elevation is associated with kidney damage and may accelerate the progression of diabetic nephropathy, also correlating with the advancement of renal insufficiency in type 2 diabetes^46^. The correlation between MCP-1 and proteinuria reinforces its role in monitoring silent kidney lesions. NGAL showed a trend toward elevation, suggesting possible usefulness in detecting acute tubular injury, although without statistical significance. Studies show that the inflammatory biomarkers MCP-1, MDA, and SYN-1 are effective in the early detection of renal dysfunctions in other parasitic infections, before changes in traditional parameters occur. MCP-1 and MDA indicate inflammation and early renal dysfunction in visceral leishmaniasis^47^. Increased SYN-1 is associated with destruction of the endothelial glycocalyx and acute renal failure in leptospirosis^38^ and shows progressive elevation in the chronic phase of Chagas disease, indicating ongoing damage. In intestinal schistosomiasis, elevated urinary MCP-1 is associated with albuminuria, and indicates silent renal inflammation despite treatment, highlighting it as a sensitive marker of renal injury^48^.

The findings of this study align with those of Oliveira et al.^47^, which reported tubuloglomerular dysfunction and evidence of renal inflammation in visceral leishmaniasis. Together, these studies underscore the importance of novel biomarkers for the early detection of kidney injury in parasitic infections. Both show that changes in distal tubule function and increased intracellular inflammatory markers can occur eben when traditional key function tests appear normal, suggesting that inflammation and tubuloglomerular dysfunction begin early in the disease course. Additionally, structural kidney damage markers like NGAL and KIM-1 have proven useful in detecting acute kidney injury (AKI) earlier than serum creatinine. Supporting this, a pilot study by Van Wolwesinkel et al.^49^ found that both NGAL and KIM-1 effectively diagnosed identified AKI in travelers with by *P. falciparum*, with urinary NGAL showing particularly strong predictive performance. These align with our study, which observed elevated NGAL levels across different phases of Chagas Disease.

NGAL levels were elevated in the chronic CD patients, especially in group G2a, which included patients without comorbidities (diabetes or systemic arterial hypertension). This suggests that NGAL elevation may reflect renal and cardiac alterations directly resulting from Chagas disease. In group G2b, where NGAL was also elevated, only three patients had hypertension, but all exhibited cardiac involvement, reinforcing this association. Although higher NGAL levels were observed in G2, an increase was also observed in the acute groups (G1a and G1b), however not statistically significant. NGAL emerges as a promising biomarker whose usefulness extends beyond kidney injury, with growing evidence supporting its relevance in various pathological conditions^22^. Although its clinical applications is still being developed, our results reinforce NGAL’s potential in transforming diagnosis of renal injury and cardiovascular diseases in CD. Our findings demonstrate that MCP-1, ANG-2, and SYN-1 are elevated early in the course of CD, even when conventional kidney function tests remain normal. This suggests that inflammation, tubulointerstitial, and endothelial dysfunction are early and common features in tropical parasitic infections. These biomarkers hold potential for detecting subtle kidney injury before clinical manifestations arise allowing for earlier intervention and improved disease management before kidney function complications.

Effective monitoring of Chagas disease in the Amazon region requires surveillance strategies adapted to the local context. While integrating Chagas disease detection, diagnosis, and follow-up into existing endemic disease programs, like as malaria, has shown promise, gaps remain in clinical follow-up after treatment and in the early detection of renal and cardiac injury. In CD infections, renal damage may progress silently and potentially worsening long-term patient prognosis. This study demonstrated that classical renal function biomarkers like creatinine and urea, underperformed in detecting early lesions associated with CD. Their lack of correlation with emerging indicators like SYN-1, ANG-2, MCP-1, and NGAL underscore the need to further develop and incorporate these novel biomarkers as sensitive tools and a means of detecting early subclinical changes in the context of CD before dysfunctions occur

### Conclusion

The study highlights that Chagas disease in the Amazon presents specific characteristics, such as the predominance of oral transmission and difficulties in accessing healthcare services, compromising early diagnosis and clinical follow-up. The use of non-traditional biomarkers, especially NGAL, SYN-1, ANG-2, and MCP-1, proved promising for the early detection of renal injuries and vascular dysfunctions associated with the disease. The elevation of these biomarkers in different clinical phases reveals the role of inflammatory and endothelial processes in the pathogenesis of renal and cardiovascular complications, often before detected by traditional diagnostic indicators. They are also reliable markers irrespective of disease stage.

Urinary MCP-1 increased in the acute pre-treatment phase, regardless of the presence of comorbidities, and significant correlations were found between ANG-2 and MCP-1 with proteinuria, as well as between ANG-2 and glomerular filtration rate. Early recognition of these alterations is crucial for specific interventions that can slow the progression of renal damage and improve patient prognosis.

These biomarkers demonstrate potential as more sensitive and individualized tools for clinical monitoring, contributing to advances in understanding pathogenesis and therapeutic management of the disease. Together with strategies like professional training, civic education and public awareness campaigns, mandatory patient follow-up, and expanded access to healthcare services, implementing the use of these biomarkers in clinical diagnosis can significantly improve the clinical outcomes of Chagas disease in the region.

### Limitations of the study

This study had several limitations. The difficulty in the longitudinal monitoring of patients, evidenced by the lack of follow-up for clinical evaluations in over 40% of cases, compromises the epidemiological and clinical assessment of disease progression in the Amazon region. Limited accessibility to healthcare services and low adherence to post-treatment follow-up hinder adequate clinical monitoring of patients with CD in the region, thus affecting the collection of complete and longitudinal data for deeper analysis. Finally, the low prevalence of chronic phase patients and those with severe cardiac involvement limited a detailed analysis of these subgroups, making it difficult to generalize findings to all disease stages.

### Perspectives and future recommendations

Develop further the non-traditional renal and endothelial biomarkers (SYN-1, ANG-2, MCP-1, and NGAL) for inclusion in the diagnosis and prognostic monitoring of early detection of renal and vascular dysfunctions in CD. Continuous monitoring strategies are essential for effective therapeutic management, especially in remote areas with limited access to healthcare services, aiming to reduce morbidity and mortality associated with the disease. Future studies with more robust longitudinal follow-up are necessary to confirm the clinical-epidemiological profile and validate the role of these biomarkers for early detection, interventions, and improved care for patients.

## Acknowledgments

The authors wish to thank patients with Chagas Disease for their participation in this study, and the public health surveillance teams. We also acknowledge the following institutions for the support they accorded: Fundação de Medicina Tropical Dr. Heitor Vieira Dourado, Amazonas Health Surveillance Foundation Dr. Rosimary Costa Pinto (FVS-RCP/AM), the Municipal Health Departments in the affected municipalities, the Pharmaceutical Bioprospecting and Clinical Biochemistry Laboratory (LBFBC) of the Federal University of Ceará.

We appreciate Victor Irungu Mwangi for the English language revision and manuscript copyediting assistance.

